# Expectation modulates the hedonic experiences of and midbrain responses to sweet flavour

**DOI:** 10.1101/2025.05.09.651338

**Authors:** Elena Mainetto, Margaret L. Westwater, Hisham Ziauddeen, Kelly M.J. Diederen, Paul C. Fletcher

**Author notes:** Please address correspondence regarding this manuscript to: Name: Paul C. Fletcher, Address: Department of Psychiatry, University of Cambridge Clifford Allbutt Building, Addenbrooke’s Hospital, Cambridge CB2 0SZ, United Kingdom. These authors contributed equally (Elena Mainetto, Margaret L. Westwater).

## Abstract

Non-nutritive sweeteners are sugar substitutes that may promote weight management by reducing an individual’s calorie intake. It is, however, unclear whether (i) sugar and non-nutritive sweetener elicit distinct orosensory responses in the human brain, and (ii) whether the neural responses to these flavours are modulated by expectancy. Addressing these questions has direct relevance to our understanding of food choice behaviour and how it may be modified in dietary interventions. We screened N=99 healthy adults to select a sample (N=27, M[SD]_age_ = 24.25[2.94] years) who reported similar perceptual experiences of sugar and sweetener, thus removing a potential confound of sensory differences, for fMRI scanning. While scanning, they received sugar- and artificially-sweetened beverages in two conditioning paradigms, which both manipulated participants’ expectation of flavour delivery: first in a probabilistic and second in a deterministic way. Participants’ ability to accurately distinguish sugar from non-nutritive sweetener depended largely on their expectations, which also significantly affected the perceived pleasantness of each flavour. Expectation altered brain responses to flavour delivery during the deterministic task only, where the (mistaken) expectation of sugar significantly increased midbrain responses to sweetener compared to when sweetener was expected. Trial-wise confidence and pleasantness ratings differentially augmented brain responses to sugar and sweetener delivery. These results highlight the importance of expectancy in both the behavioural and neural encoding of sweet flavour, particularly in the context of unreliable sensory information. The expectation of sugar appears to increase the subjective value of noncaloric sweetener, which may result from flavour-nutrient conditioning that preferentially reinforcers sugar.

## 1. Introduction

Global trends in overweight and obesity have driven considerable research into the reinforcing properties of highly-palatable, energy-dense foods, which can contribute to difficulties with weight management. While such foods typically have both high fat and high sugar content, sugar remains a common target of policy and intervention efforts surrounding obesity (1,2). Some efforts promote non-nutritive sweeteners, such as saccharin, sucralose, or aspartame, as a sugar-sweetened beverage substitute that reduces excessive caloric intake but maintains a similarly rewarding flavour. However, little is known about how sugar and non-nutritive sweetener consumption are encoded in the human brain. Advancing knowledge of this mechanism will have key implications for our understanding of food choice behaviour and appropriate dietary interventions particularly since empirical support for non-nutritive sweeteners as a means of weight management remains inconclusive (3–5).

Findings from preclinical neuroscience have identified distinct neural pathways that signal the hedonic and nutritive value of sugar, which broadly correspond to the ingestive and post-ingestive effects of consumption. The hedonic value of sweet flavour arises from orosensory inputs (e.g., taste, olfaction) that depolarize taste receptors, which project first to the nucleus tractus solitarius and subsequently to the thalamic ventroposterior medial nucleus and primary gustatory cortex (i.e., anterior insula and frontal operculum (6)). Subregions of the insular cortex then project to reward-related brain areas, including the ventromedial prefrontal cortex and ventral striatum (7). Indeed, both non-nutritive sweetener and sugar augment dopamine release from the nucleus accumbens, even in sham-fed animals (8,9), suggesting that ventral striatal areas selectively encode the reinforcing effects of sweetness. In contrast, the nutritive value of sugar is signalled by melanin-concentrating hormone expressing neurons in the lateral hypothalamus that project to the dorsal striatum (10). This central pathway also receives peripheral input from hepatic portal vein sensors via the gut-brain axis (10).

While many of the aforementioned studies exposed animals to both non-nutritive and caloric sweeteners, comparatively few studies have investigated neural responses to these tastants in humans. Initial findings suggested that both sucrose and sucralose elicit gustatory cortex responses, but only sucrose delivery augmented ventral striatal activation (11). Ventral striatal responses to caloric sweetener may be modulated by sensory-specific satiety, whereas greater amygdala activation following caloric versus non-caloric sweetener delivery is reportedly independent of satiety (12). Although these effects could suggest that individuals find non-nutritive sweeteners less rewarding, the limited sample size (n=10) and correspondingly low statistical power of these studies warrants caution. Indeed, meta-analytic findings indicate robust anterior insula and frontal operculum responses to sweeteners, whereas neither caloric nor non-nutritive sweeteners reliably elicit ventral striatal responses. However, aligning with preclinical findings, dorsal striatal activation increases following caloric sweetener delivery (13).

Another challenge to human neuroscience studies of artificial sweeteners relates to potential differences in the expected value of these flavours relative to sugar. Non-nutritive sweeteners have an intense sweetness that is tens or hundreds of times that of sugar, and differences in their affinity for “sweet” receptors (T1R2 and T1R3 (14)) on the tongue may also contribute to qualitatively different perceptions of these tastants. Moreover, non-caloric sweeteners also bind to T2 receptors that encode bitter taste, often making them easily distinguishable from caloric sweeteners and less pleasant. These orosensory properties may lead one to anticipate non-nutritive sweeteners to be less palatable, thereby reducing their reward value through reinforcement learning (15,16)). Midbrain dopaminergic prediction error (PE) signals, which code the difference between expected and actual value, underlie reinforcement learning, where both the sign and magnitude of a PE contribute to the reward value of a given outcome. As such, the apparently less rewarding effects of non-nutritive sweeteners relative to sugar may relate to differences in not only their associated PEs and their intrinsic hedonic properties but also learning-based expectation.

Given the uncertainties about both the potential differences in perceptual experiences between sugar and non-nutritive sweetener, as well as the likely influences of reward expectancies, we sought to characterise the neural responses to the delivery of either sugar- or non-nutritive sweetener sweetened beverages. Critically, we examined these processes within participants who had been shown, through pre-screening, to be unable to accurately distinguish these flavours, thus ensuring that we could manipulate expectancies independently from perceptual experience of each stimulus. We used trial-specific cueing to engender varyingly strong expectations about whether the ensuing liquid delivery would contain sugar or sweetener. In the first task, expectations were probabilistic (75%, 50% or 25% chance of a sugar-sweetened beverage), and in the subsequent paradigm, strong expectations were generated by a deterministic design that informed participants that the next flavour would definitely be sugar or non-nutritive sweetener. We examined the impact of these expectations (and their violation) on task performance and neural responses to sugar and sweetener across both gustatory and reward networks. For the probabilistic paradigm, we anticipated that participants would show improved accuracy for high-probability flavours, and accuracy would be reduced following low-probability sugar or sweetener delivery, relative to chance. We predicted that expectation would modulate pleasantness ratings during the deterministic paradigm, such that the expectation of sugar would be associated with greater pleasantness. At the neural level, delivery of both sugar and non-nutritive sweetener was predicted to elicit similar neural responses across gustatory regions, aligning with participants’ poor discriminability of the flavours. However, we anticipated that the expectation of sugar would be associated with increased activation of putative reward regions, whereas the expectation of sweetener would manifest as an attenuated neural response, irrespective of the flavour delivered. Finally, exploratory analyses examined the parametric effect of confidence and pleasantness ratings on brain responses to each flavour for the probabilistic and deterministic tasks, respectively.

## 2. Results

Ninety-nine healthy volunteers attended a screening visit, and twenty-seven participants (N=19 female, M[SD] age = 24.25[2.94] years) who could not accurately discriminate between sugar- and artificially-sweetened beverages were invited to undergo fMRI scanning in Cambridge, UK (see **Methods**). During the scan, participants performed probabilistic and deterministic conditioning tasks, which each manipulated expectations regarding whether there would be a subsequent delivery of sugar or artificial sweetener solution (see **Figure 1A-C** & **Methods**). On each trial, participants provided confidence (i.e., about which taste they received) and pleasantness ratings following beverage delivery for probabilistic and deterministic phases, respectively. Hunger, thirst and liking ratings for each flavour were recorded at three time points throughout the scan. Neuroimaging data were pre-processed using SPM12 software (Matlab 2016b) with 6mm spatial smoothing, and first-level statistical analyses modelled the effect of expectation and flavour perception across probabilistic and deterministic phases (see **Methods**). Group-level analyses were initially constrained to *a priori* regions of interest (ROIs) that have previously been implicated in reward expectation and flavour perception (see **Figure 2**); however, these were succeeded by an exploratory whole-brain analyses for each contrast of interest.

**Figure 1.**
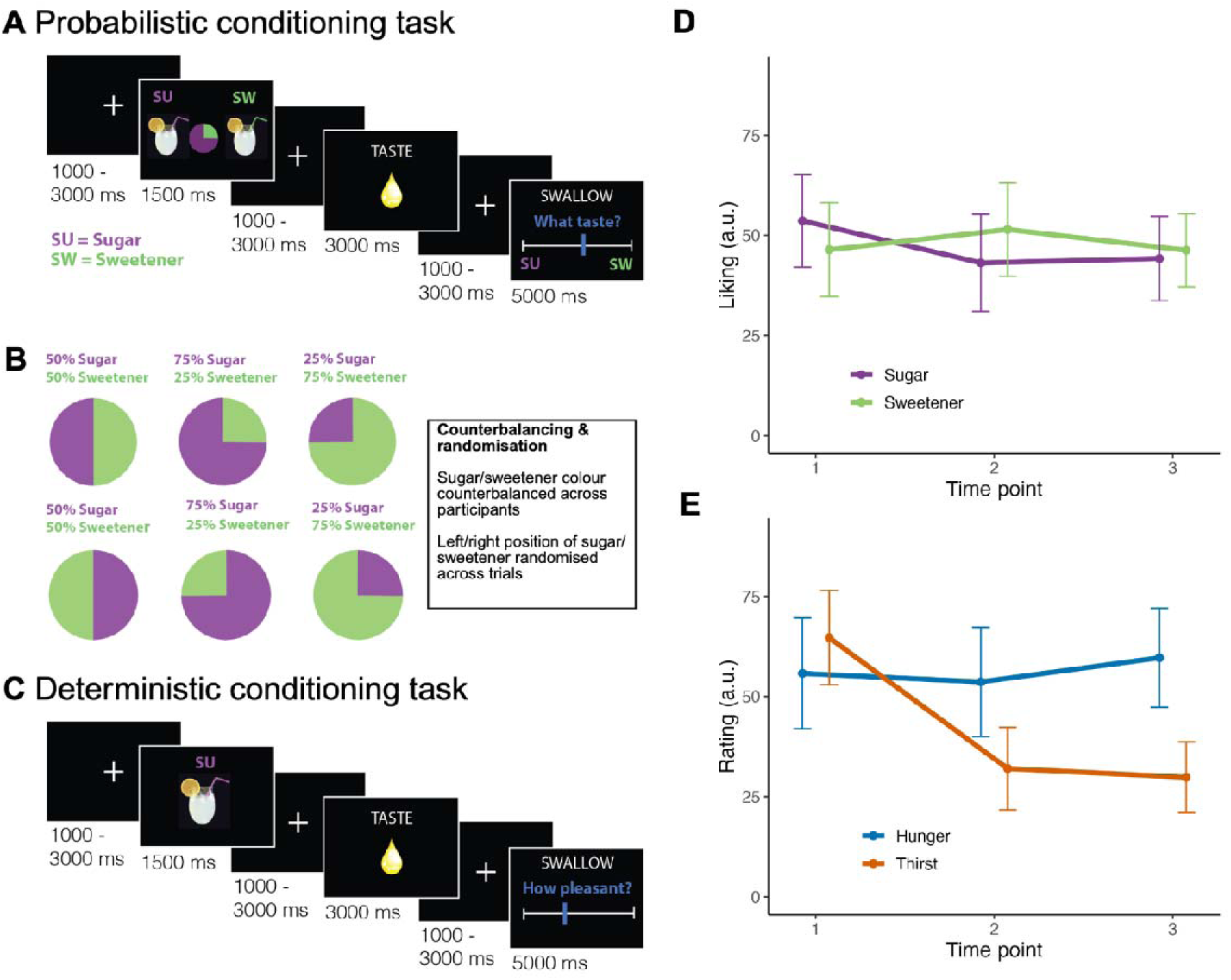
Probabilistic and deterministic conditioning task trial structure and internal state ratings. **A)** Probabilistic conditioning task trial structure. A cue indicating the probability of sugar-sweetened lemonade and non-nutritive sweetener lemonade was presented prior to flavour stimulus delivery. Participants were instructed to hold the liquid in their mouth until the “SWALLOW” cue was presented, at which time they rated which flavour they received. **B)** The associated colour (green or purple) for each flavour was counterbalanced across participants, and the position of each flavour (left or right) on the slider was randomised across trials. **C)** Deterministic conditioning task trial structure. On each trial, a cue depicting either sugar-sweetened lemonade or non-nutritive sweetener lemonade was followed by delivery of the cued liquid. However, cues were covertly violated on 50% of trials (except for the neutral water rinse). Participants were then instructed to swallow the liquid and rate the pleasantness of the flavour. **D)** Liking ratings did not differ significantly between sugar and non-nutritive sweetener, and they remained stable during the experiment. **E)** Internal state ratings throughout the MRI scan showed that participants’ thirst significantly decreased while hunger remained unchanged. Created with Biorender.com.

**Figure 2.**
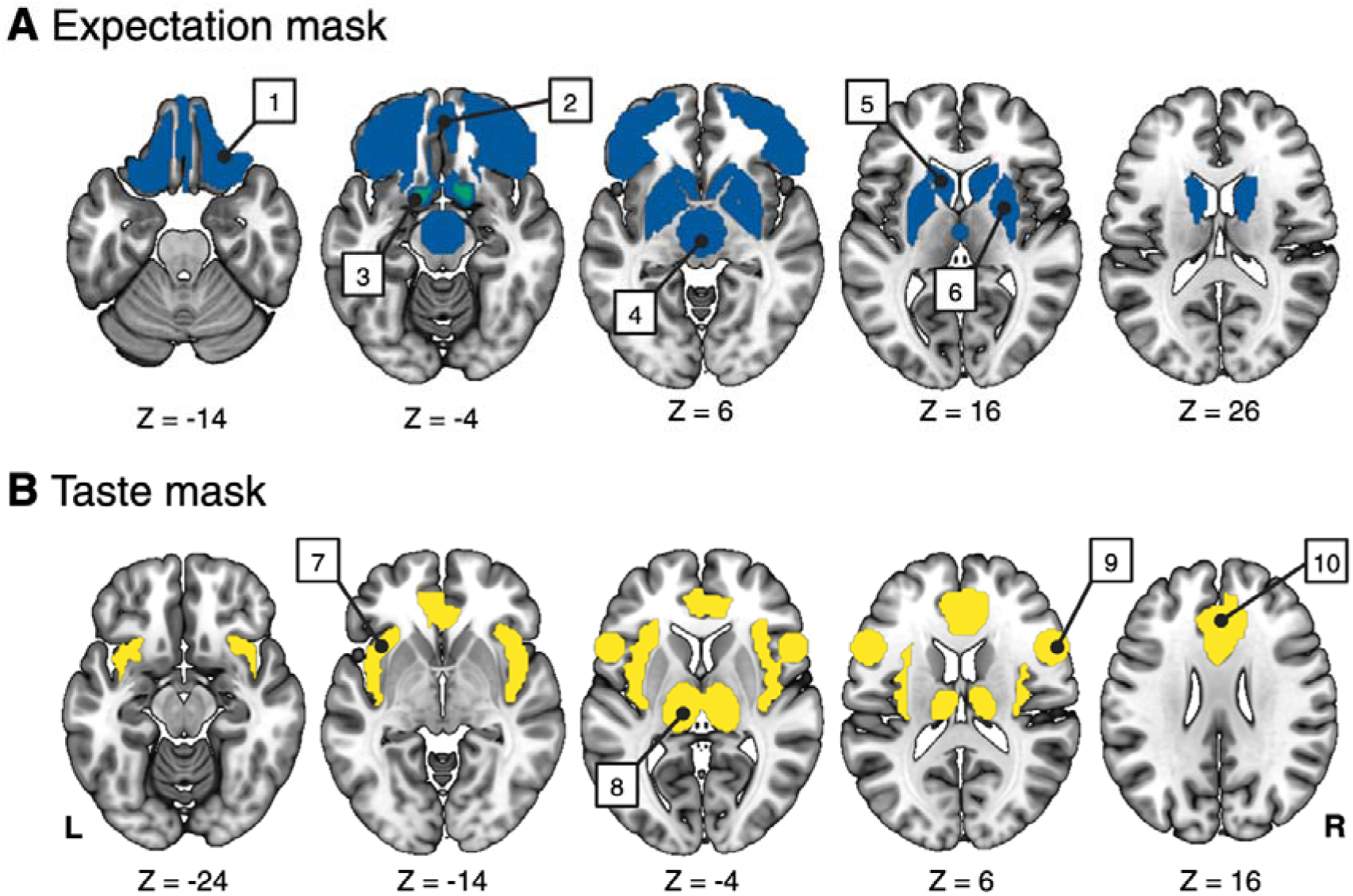
Regions of interest in taste and expectation masks. **A)** The expectation mask was composed of bilateral basal ganglia and ventral frontal regions: 1 = orbitofrontal cortex, 2 = ventromedial prefrontal cortex, 3 = nucleus accumbens, 4 = midbrain, 5 = caudate, 6 = putamen. **A)** The taste mask included bilateral regions that have been functionally implicated in flavour perception: 7 = insula, 8 = thalamus, 9 = frontal operculum, and 10 = anterior cingulate cortex. Created with Biorender.com.

### 2.1 Behavioural results

Behavioural data were analysed with linear mixed-effects models (LMMs) using the R package *nlme* (17). Liking ratings did not differ significantly between flavours (p = 0.80), and they did not change from baseline (**Figure 1D**). A flavour-by-time interaction effect was nonsignificant, indicating that liking ratings for each flavour remained similar for the duration of the experiment (**Figure 1E**). Thirst ratings decreased across all time points, whereas hunger levels remained stable (see **Supplementary Material**).

LMMs for the probabilistic task examined the effect of task conditions on flavour identification accuracy and confidence, whereas deterministic task models examined these effects on pleasantness (see **Methods**). On the probabilistic task, the main effect of flavour type on accuracy was nonsignificant, indicating that participants were unable to consistently distinguish the flavours. Moreover, a significant main effect of probability level indicated that participants were nominally less accurate for unexpected trials relative to chance (B(SE) = - 5.74(2.58), p = 0.032), and they were more accurate on trials when the expected flavour was delivered as compared to chance (B(SE) = 9.22(2.59), p = 0.001; **Figure 3A**). This would suggest that the task successfully manipulated participants expectations regarding flavour delivery. All interaction effects on accuracy were nonsignificant. The main and interaction effects of flavour type and probability level were not significantly related to confidence ratings (all p’s >.14).

**Figure 3.**
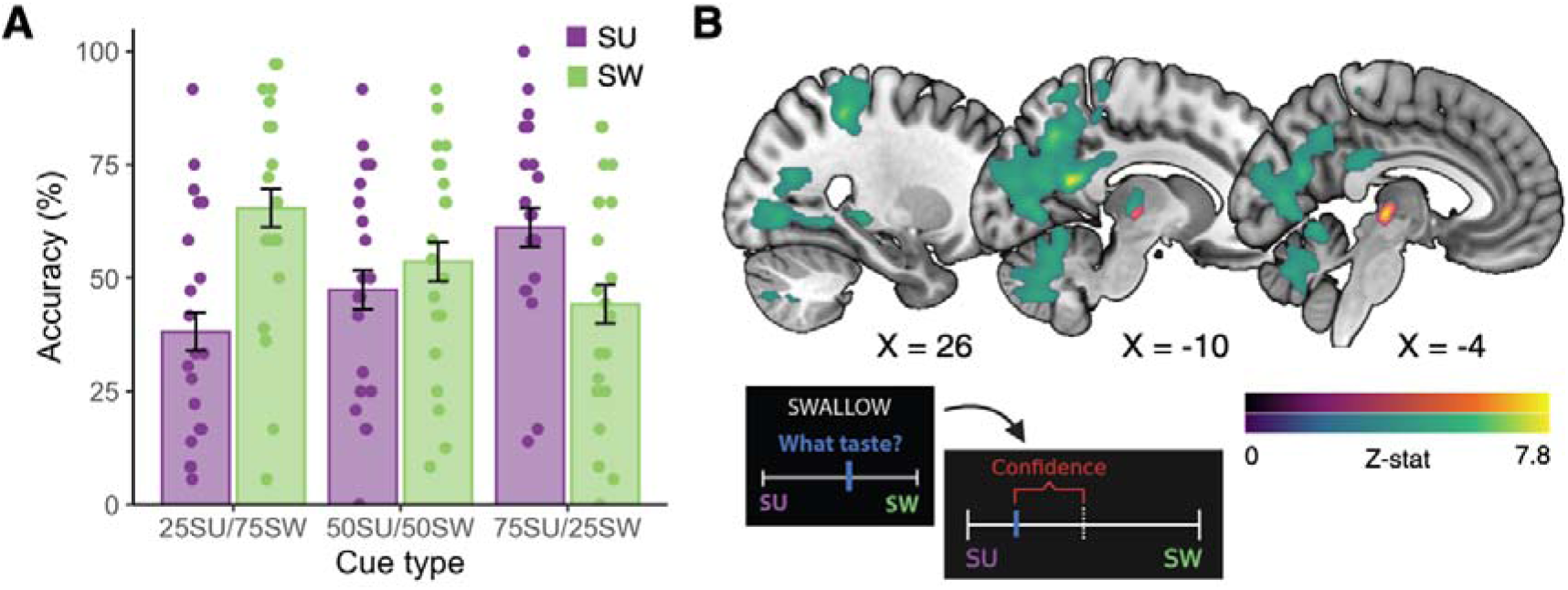
Probabilistic task performance and neural encoding of confidence. **A)** The probability of flavour delivery significantly affected accuracy, which was greater for expected flavours and nominally lower for unexpected flavours, relative to chance. Accuracy did not differ across flavour types. Colours depict the flavour delivered. Error bars = 95% confidence interval. **B)** The parametric effect of trial-by-trial confidence differed between sugar and non-nutritive sweetener trials in both our a priori ROI analyses (purple/orange colour bar, voxel-wise p < .001, small volume corrected p < .05) and whole-brain analyses (blue/green colour bar, voxel-wise p < .001, FEW-corrected cluster probability p < .05). For details on cluster size, coordinates, and associated test statistics of these one-tailed t-tests, see Tables 2 & S1. Confidence was calculated from taste identification ratings as the absolute difference of the response (blue) from the midpoint (dashed line), which was divided by the range and scaled by 100.

LMMs for the deterministic task showed that the main effects of flavour and violation of expectation on pleasantness ratings were nonsignificant (p-value’s > 0.05); however, a significant flavour-by-violation (of expectation) interaction effect indicated that pleasantness of the non-nutritive versus sugar-sweetened lemonade was significantly greater on unexpected versus expected trials (B(SE) = 14.06(1.74), p < 0.001). That is, participants found non-nutritive sweetener to be more pleasant when they were expecting sugar than when they were expecting sweetener, and the opposite was observed for sugar (**Figure 4A**). Inclusion of the interaction term significantly improved model fit (*χ*^2^(_1_) = 64.56, p < 0.001).

**Figure 4.**
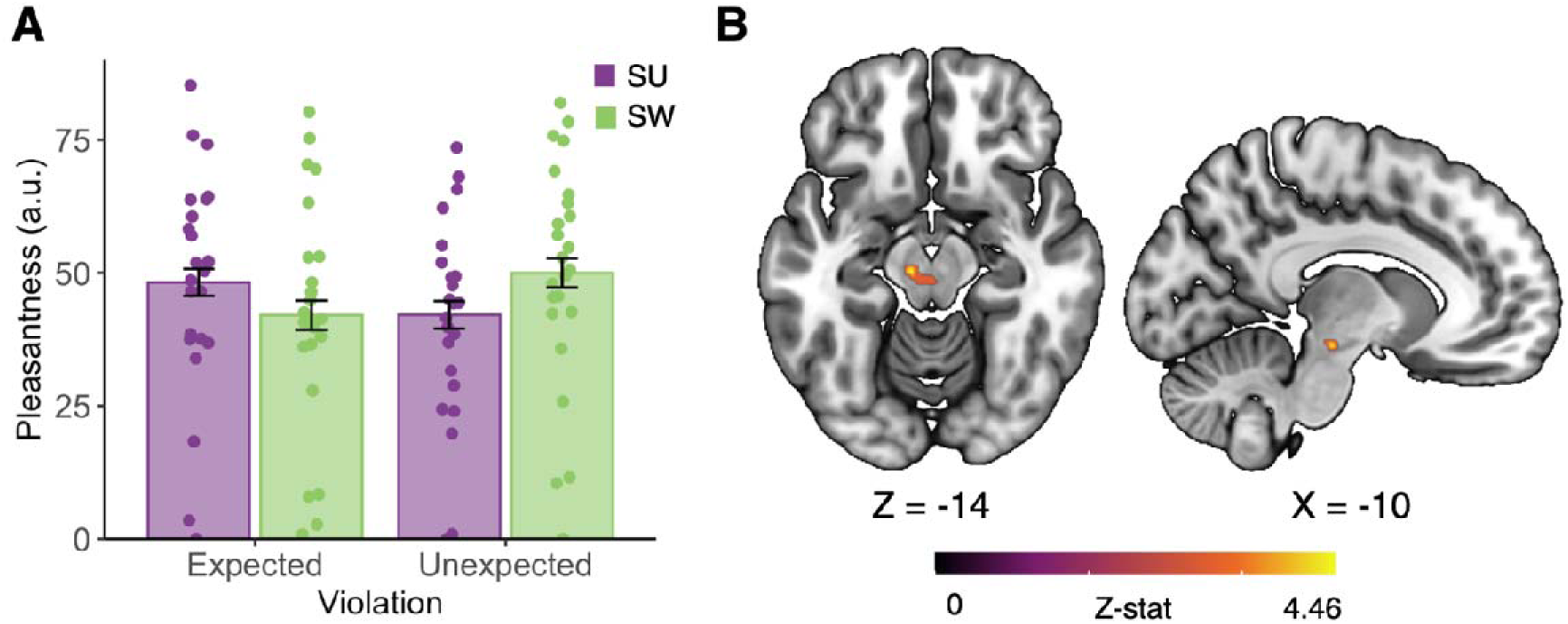
Expectation of sugar augments pleasantness ratings and midbrain responses to non-nutritive sweetener. **A)** A flavour-by-violation interaction indicated that participants found unexpected non-nutritive sweetener to be more pleasant than unexpected sugar. That is, participants rated non-nutritive sweetener as more pleasant when they thought they had received sugar (and vice versa for sugar). Colours depict the flavour delivered. Error bars = 95% confidence interval. **B)** This effect corresponded with increased left midbrain activation for unexpected versus expected non-nutritive sweetener delivery (t-stat = 5.78, peak = .024, 46 voxels). Created with Biorender.com.

### 2.2 Neuroimaging results: Probabilistic task

During the probabilistic conditioning task, participants’ expectations were manipulated by presenting visual cues depicting varying probability levels (25%, 50%, 75%) of either sugar- or artificially-sweetened beverage delivery. We were therefore able to examine brain responses to high-probability versus uncertain sugar (75% sugar cue vs 50% sugar cue) and non-nutritive sweetener (75% sweetener cue vs 50% sweetener cue), as well as low-probability versus uncertain sugar (25% sugar cue vs 50% sugar cue) and sweetener (25% sweetener cue vs 50% sweetener cue; **Figure 1A & B**). However, it should be noted that the high-probability cues for one flavour also conveyed a low probability of the other. Our primary ROI analyses indicated that brain responses across striatal, medial frontal, orbitofrontal and midbrain regions were not significantly different across probability levels for both sugar and non-nutritive sweetener. The main effect of probability level was largely nonsignificant at the whole-brain level; however, left superior occipital, lingual gyrus and cerebellum activation was greater for uncertain (50% probability) cues as compared to the other probability levels (**Table S1**).

When examining neural responses to flavour delivery, sugar-sweetened beverages elicited increased left putamen activation, whereas artificially sweetened beverages increased brain responses in several regions that have established roles in gustatory and reward processing (see **Table 2, Figure S1**). The main effect of flavour type across all probability levels was not significant, suggesting markedly similar brain responses to sugar and non-nutritive sweetener delivery, which mirrored participants’ behavioural ratings of these flavours. However, exploratory whole-brain analyses indicated that, on maximally uncertain trials (50%), left superior parietal activation was increased following non-nutritive sweetener versus sugar delivery (**Table S1)**.

**Table 1.**
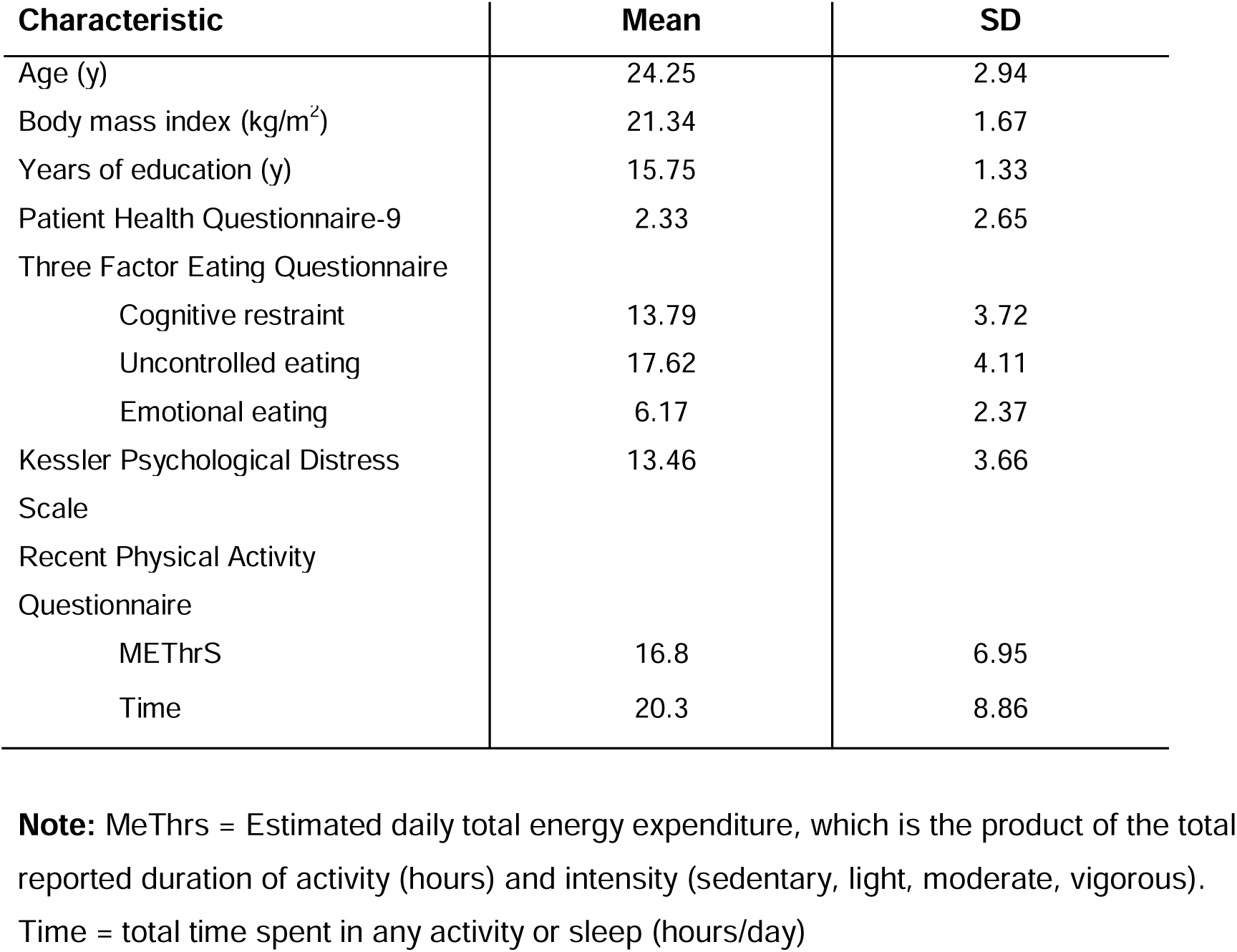
Demographic information.

**Table 2.**
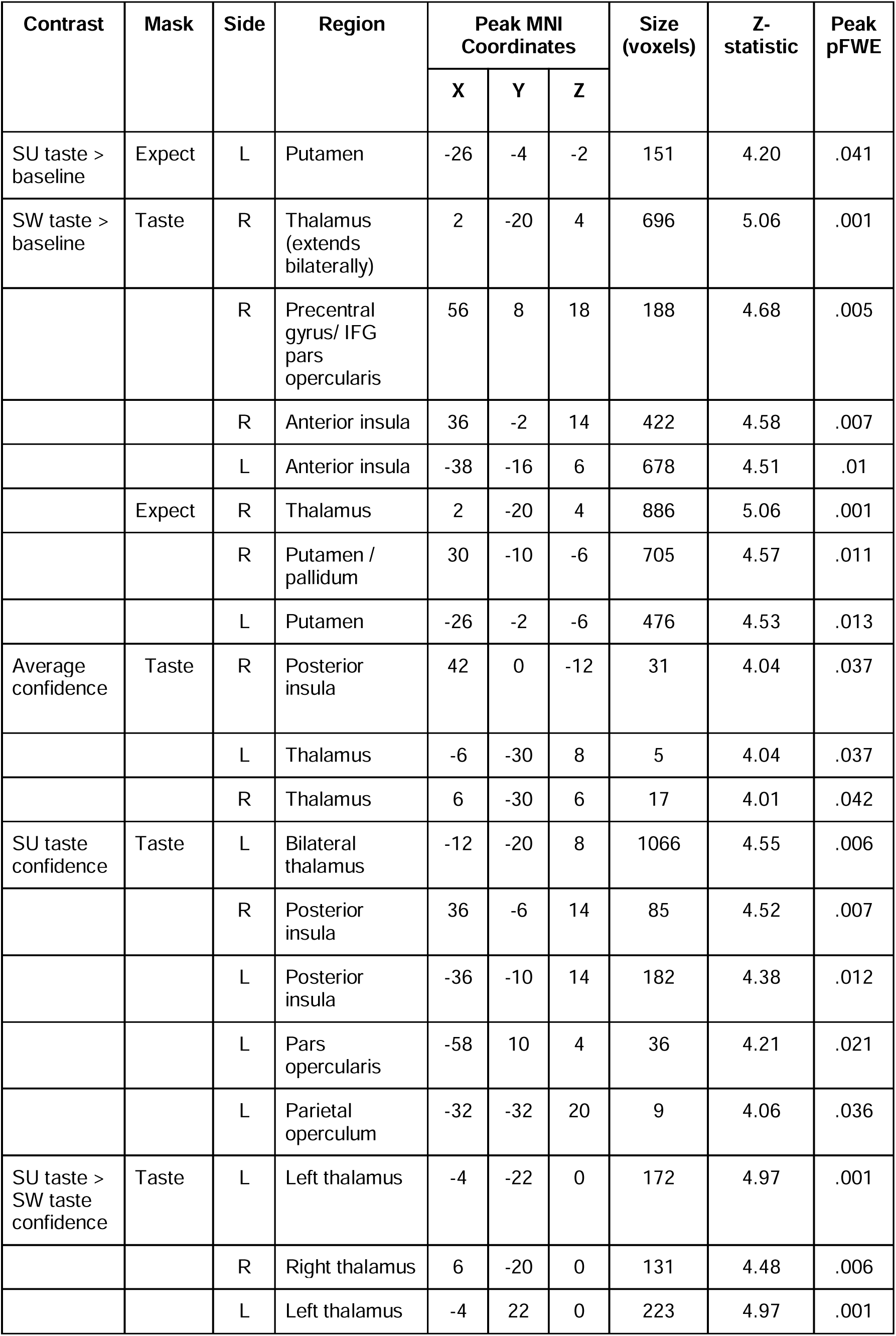
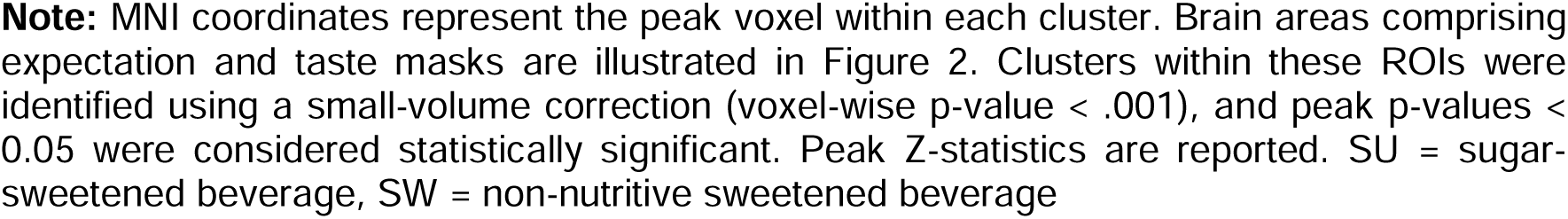
ROI brain responses during the probabilistic conditioning task.

| Contrast | Mask | Side | Region | Peak MNI Coordinates |  |  | Size (voxels) | Z-statistic | Peak pFWE |
| --- | --- | --- | --- | --- | --- | --- | --- | --- | --- |
|  |  |  |  | X | Y | Z |  |  |  |
| SU taste > baseline | Expect | L | Putamen | -26 | -4 | -2 | 151 | 4.20 | .041 |
| SW taste > baseline | Taste | R | Thalamus (extends bilaterally) | 2 | -20 | 4 | 696 | 5.06 | .001 |
|  |  | R | Precentral gyrus/ IFG pars opercularis | 56 | 8 | 18 | 188 | 4.68 | .005 |
|  |  | R | Anterior insula | 36 | -2 | 14 | 422 | 4.58 | .007 |
|  |  | L | Anterior insula | -38 | -16 | 6 | 678 | 4.51 | .01 |
|  | Expect | R | Thalamus | 2 | -20 | 4 | 886 | 5.06 | .001 |
|  |  | R | Putamen / pallidum | 30 | -10 | -6 | 705 | 4.57 | .011 |
|  |  | L | Putamen | -26 | -2 | -6 | 476 | 4.53 | .013 |
| Average confidence | Taste | R | Posterior insula | 42 | 0 | -12 | 31 | 4.04 | .037 |
|  |  | L | Thalamus | -6 | -30 | 8 | 5 | 4.04 | .037 |
|  |  | R | Thalamus | 6 | -30 | 6 | 17 | 4.01 | .042 |
| SU taste confidence | Taste | L | Bilateral thalamus | -12 | -20 | 8 | 1066 | 4.55 | .006 |
|  |  | R | Posterior insula | 36 | -6 | 14 | 85 | 4.52 | .007 |
|  |  | L | Posterior insula | -36 | -10 | 14 | 182 | 4.38 | .012 |
|  |  | L | Pars opercularis | -58 | 10 | 4 | 36 | 4.21 | .021 |
|  |  | L | Parietal operculum | -32 | -32 | 20 | 9 | 4.06 | .036 |
| SU taste > SW taste confidence | Taste | L | Left thalamus | -4 | -22 | 0 | 172 | 4.97 | .001 |
|  |  | R | Right thalamus | 6 | -20 | 0 | 131 | 4.48 | .006 |
|  |  | L | Left thalamus | -4 | 22 | 0 | 223 | 4.97 | .001 |

**Note:** MNI coordinates represent the peak voxel within each cluster. Brain areas comprising expectation and taste masks are illustrated in Figure 2. Clusters within these ROIs were identified using a small-volume correction (voxel-wise p-value < .001), and peak p-values < 0.05 were considered statistically significant. Peak Z-statistics are reported. SU = sugar-sweetened beverage, SW = non-nutritive sweetened beverage

The parametric effect of trial wise confidence across all trial types was associated with greater activation in primary sensory areas, including the bilateral thalamus and right posterior insula (ROI analyses; **Table 2**). Moreover, as confidence increased, bilateral thalamic responses following sugar versus non-nutritive sweetener delivery also increased; however, there were no significant differences across probability levels. At the whole-brain level, increasing confidence was related to increased activation across three large clusters, spanning hippocampal, parahippocampal, primary sensory, and gustatory regions (**Table S1 & Figure S3**). Moreover, the parametric effect of confidence significantly increased activation in one large cluster, spanning superior parietal, inferior parietal, and thalamic regions, following sugar delivery relative to non-nutritive sweetener (**Figure 3B & Table S1**). Perceptual decision-making experiments have implicated hippocampal and parietal regions in the estimation of choice uncertainty (18,19), and our findings suggest that this circuit may also code confidence estimates in relation to gustatory stimuli.

### 2.3 Neuroimaging results: Deterministic task

Following the probabilistic task, participants performed a deterministic conditioning paradigm in which they viewed cues for either sugar, non-nutritive sweetener or water beverages, and these were followed by a 0.9mL delivery of the cued liquid. However, the cues were covertly violated on 50% of sugar and sweetener trials (that is, a sugar cue was succeeded by sweetener delivery and vice versa; see **Methods**). As in the probabilistic task, delivery of both sugar- and artificially sweetened liquids was related to increased activation of gustatory and primary sensory regions relative to the implicit baseline (see **Table 3**). Although brain responses did not differ across unexpected versus expected sugar trials, we observed increased midbrain activation on unexpected versus expected sweetener trials in our ROI analysis (**Figure 4B**; MNI_X,Y,Z_ = -10, -22, -14, Z = 4.46, p = 0.024, small-volume corrected). This may suggest that the ventral striatum preferentially encodes the value of sweet flavour when nutritive value is expected.

**Table 3.** ROI brain responses during the deterministic conditioning task.

| Contrast | Mask | Side | Region | Peak MNI Coordinates |  |  | Size (voxels) | Z-statistic | Peak pFWE |
| --- | --- | --- | --- | --- | --- | --- | --- | --- | --- |
|  |  |  |  | X | Y | Z |  |  |  |
| Unexpected SW > Expected SW taste | Expect | L | Midbrain, substantia nigra pars compacta | -10 | -22 | -14 | 46 | 4.46 | .024 |
| SU taste > baseline | Expect | L | Pallidum, putamen | -26 | -14 | -4 | 89 | 4.25 | .022 |
|  | Taste | L | Posterior insula | -36 | -12 | 14 | 184 | 4.31 | .012 |
|  |  | R | Posterior insula | 36 | -6 | -14 | 125 | 4.27 | .015 |
|  |  | R | Thalamus | 12 | -16 | 6 | 186 | 3.98 | .041 |
|  |  | L | Thalamus | -10 | -20 | 0 | 152 | 3.93 | .048 |
| SW taste > baseline | Taste | L | Posterior insula | -36 | -12 | 14 | 139 | 4.18 | .021 |
|  |  | R | Posterior insula | 36 | -6 | 12 | 91 | 4.12 | .025 |
| Pleasantness SU taste | Taste | L | Posterior insula, central operculum | -38 | -12 | 18 | 164 | 4.70 | .004 |
|  |  | R | Thalamus | 12 | -14 | 6 | 175 | 4.21 | .024 |
|  |  | R | Posterior insula, central operculum | 40 | -8 | 22 | 88 | 4.11 | .035 |
|  |  | L | Parietal operculum, posterior insula | -32 | -32 | 20 | 23 | 4.09 | .037 |

**Note:** MNI coordinates represent the peak voxel within each cluster. Brain areas comprising expectation and taste masks are illustrated in Figure 2. Clusters within these ROIs were identified using a small-volume correction (voxel-wise $p < .001$ ); peak $p$ -values $< 0.05$ were considered statistically significant. Peak Z-statistics are reported. SU = sugar-sweetened beverage, SW = non-nutritive sweetened beverage

Since our behavioural results suggested that expectation modulates the subjective pleasantness of sweet flavour, we interrogated the parametric effect of pleasantness on brain responses following liquid delivery. As pleasantness increased, brain responses to sugar also increased in the bilateral insula and right thalamus, and this effect did not differ across expected versus unexpected trials (ROI analyses; **Table 3**). Exploratory whole-brain analyses indicated that these effects extended to two large clusters, which encompassed several regions known to be associated with sweet flavour perception and interoception (e.g., postcentral gyrus, basal ganglia, brainstem nuclei, cerebellum; see **Table S2 & Figure S4**). Although pleasantness ratings did not alter brain responses to non-nutritive sweetener, the parametric effect of pleasantness differentially modulated lateral occipital responses to sugar versus non-nutritive sweetener (**Table S2**). This effect did not differ by expectation.

## 3. Discussion

The present study implemented two conditioning paradigms to characterise how expectations of sugar or non-nutritive sweetener affect the neural coding of sweet flavour. Critically, to control for differences in the sensory experience of each flavour, we assessed these processes in healthy adults who were unable to reliably distinguish sugar- and non-nutritive sweetened beverages. Analysis of both behavioural and neuroimaging data yielded three key results. First, in a probabilistic conditioning task, probability level significantly affected flavour identification accuracy, and participant’s confidence in their choice increased brain responses across primary sensory, hippocampal and parietal clusters. Second, during the deterministic task, expectation had opposing effects on the pleasantness of non-nutritive sweetener and sugar, where the expectation of sugar was associated with enhanced subjected pleasantness when participants unknowingly received non-nutritive sweetener. Increased pleasantness ratings augmented brain responses to sugar, but not sweetener, across a putative gustatory network. Finally, the expectation of sugar significantly augmented midbrain responses to covert sweetener delivery, suggesting that the expectation of nutritive value accounts, at least in part, for the distinct reinforcing effects of caloric and non-caloric sweeteners.

Findings from our probabilistic conditioning paradigm indicated that participants relied primarily on predictive cues to distinguish sugar and non-nutritive sweetener. When examining the main effects of flavour and probability level on accuracy, we found that only the predictive information significantly altered accuracy, where participants had improved performance for expected flavours and nominally poorer accuracy for unexpected ones. However, these cues were associated neither with differences in mesolimbic activation nor with participants’ confidence in their choices. The latter would suggest that participants overweighted gustatory information relative to imprecise predictive information when generating their confidence estimates. Indeed, the parametric effect of confidence was related to increased thalamic and insula activation following flavour delivery in our ROI analyses, as well as greater bilateral (para)hippocampal activation in our exploratory whole-brain analysis. Although meta-analyses support a role for the parahippocampal gyrus in retrospective confidence judgements (20,21), the present findings, to our knowledge, are the first to implicate this region in confidence estimation during gustatory decision-making. We also identified clusters in the posterior thalamus, superior parietal cortex, and occipitoparietal junction that preferentially tracked confidence following sugar versus non-nutritive sweetener delivery. Several lines of evidence suggest that parietal cortex simultaneously accumulates evidence for and codes confidence in perceptual decisions (19,22), particularly under conditions of uncertainty. Our findings broadly align with this interpretation, yet it remains unclear as to why such an effect would be specific to sugar delivery.

The deterministic conditioning paradigm further demonstrated that predictive information modulates the pleasantness of sweet flavour, suggesting that the expected value of sugar and sweetener can override their sensory properties. The direction of this effect differed between sugar and non-nutritive sweetener, where participants rated unexpected sweetener as more pleasant than expected sweetener and vice versa for sugar. That is, when sweetener delivery was preceded by sugar cues, participants rated the flavour as more palatable than when they expected sweetener. Similarly, participants found sweetener-cued sugar delivery to be less palatable than expected sugar. Moreover, the parametric effect of pleasantness was associated with greater activation across a putative gustatory network (e.g., thalamus, posterior insula, postcentral gyrus, and basal ganglia) following sugar – but not sweetener – delivery. Several lines of evidence support a conserved preference for caloric versus non-caloric sweetener across species (23,24), which likely results from post-ingestive signals (e.g., via hepatic portal vein glucose sensing, glucose metabolism) that reinforce its nutritive value in the brain (25–27). Such flavour-nutrient conditioning would augment the expected value of sugar relative to sweetener, and indeed, our results suggest that expectations surrounding the nutritive value of a sweet flavour significantly alter its palatability.

Intriguingly, we observed greater midbrain activation for unexpected versus expected sweetener delivery, yet there were no differences for unexpected sugar delivery. This may be understood within a predictive processing framework, in which unreliable sensory information has tipped the balance in favour of more precise predictive information, and consequently, the expectation of nutritive value from a sweet flavour increases its subjective value. Preclinical models indicate that the reinforcing effects of sugar are largely mediated by dorsal striatal circuitry, and peak activation of our midbrain cluster fell within the left substantia nigra (SN), which projects to the dorsolateral striatum via the nigrostriatal pathway (7). Activation of dopaminergic D1 neurons within the dorsal striatum-SN pathway has been shown to elicit calorie-seeking in rodents, even when sugar is paired with an aversive flavour (8). These effects may relate to the broader functions of the SN, which include both sensing gut-induced reward from right-lateralized vagal afferents (28) and estimation of action-based value to support goal-directed behaviour (29). Although we cannot exclude the possibility that our cluster spanned additional midbrain nuclei, our findings might suggest that the SN encodes the expectation of sugar-associated calories, which may, in turn, facilitate action selection to obtain a calorific reward.

Despite having notable strengths, several limitations of this study should be considered. First, we recruited a modest sample size of individuals who could not reliably distinguish between sugar- and non-nutritive-sweetened beverages, and it remains unclear if our findings will generalise to the broader population. Encouragingly, previous studies have found that individuals poorly discriminate between similar flavours (30), with accuracy rates of only 66% when identifying an artificially sweetened beverage (31). This may suggest that our sample reflects a substantial proportion of the general population, but replication efforts in larger samples are warranted. Second, our probabilistic task used cue stimuli that presented reciprocal probabilities for each flavour (that is, the high probability cue for sugar conveyed the same information as the low probability cue for sweetener). As such, neural responses to expected and unexpected flavours could not be fully dissociated from one another. Use of these cues increased the efficiency of our event-related paradigm, yet future studies may wish to include cues that depict probabilities for only one flavour at a time. Finally, our sample was restricted to lean volunteers, which limited our ability to examine neural responses to sweet flavour across the full BMI range. Alterations in putative sensory and reward circuitry have been identified in individuals with obesity (32–35), and our findings may guide future efforts to delineate the specific role of reward expectation in those struggling with elevated BMI and overeating.

Taken together, our findings advance understanding of the neurocognitive mechanisms that underlie the rewarding effects of sweet flavour, which are largely driven by expectation when sensory information is unreliable. The reinforcing effects of sugar appeared to trump those of non-nutritive sweetener, and this likely reflects flavour-nutrient conditioning and the evolutionarily conserved drive for calories. These results underscore the role of expectancies in guiding food choice behaviour, which may open new avenues for future dietary interventions.

## 4. Methods

### 4.1 Participants

Ninety-nine (n=54 female, M[SD] age = 23.75[3.66] years) participants were recruited from posted advertisements, university mailing lists, and social media. Eligible volunteers were healthy adults between 18 and 40 years of age with a body mass index (BMI) between 18.5 and 24.9 kg/m^2^. Exclusion criteria included: the presence of a significant medical or psychiatric history, contraindications to MRI scanning (e.g., pregnancy, some metallic implants), being left-handed, current dieting, or having English language or communication difficulties that would impede understanding of the study procedures.

### 4.2 Study procedures

Potential volunteers attended a 30-minute screening session, which involved a gustatory task (described below) that assessed participants’ ability to discriminate between the sugar and sweetener version of the same beverage, Sainsbury’s Classic Lemonade and Sainsbury’s Diet Lemonade. Participants refrained from eating or drinking anything except water for 1 hour prior to the screening. The two lemonade products had the same composition apart from the sweetening agent. Critically, the classic lemonade did not contain non-nutritive sweetener in addition to sugar. As we sought to examine the effect of expectancies on gustatory perception, we limited our sample to participants who were relatively unable to discriminate between the two liquids. Individuals with a discrimination accuracy below 50% or above 60% were excluded, as accuracy values in these ranges would reflect poor task engagement or high discrimination between the two liquids, respectively. Although these exclusion criteria were disclosed to ineligible volunteers, eligible participants were not informed of the reason for their selection. Of the screening sample, 27 participants (n=19 female) were invited to the scanning session.

MRI scan sessions began at approximately 9.00 hours, lasting 90 minutes. Participants were asked to refrain from eating or drinking anything (except water) 8h prior to the study session, and they provided internal state ratings (hunger, thirst) and liking ratings for each liquid using a Visual Analog Scale (*0 = Not at all*, *100 = Extremely*) at the beginning and end of scanning. During each scan, participants completed probabilistic and deterministic conditioning tasks, which lasted 70 minutes in total. In each task, SU and SW liquids were paired with one of two colours (green or purple). The colour association was randomised and counterbalanced across participants.

All the participants provided signed, informed consent at the beginning of the screening session, and they were compensated £60 for their time. Participants who only completed the screening visits received £10. The study was approved by the ethics committee of the University of Cambridge, Cambridge, United Kingdom (Ref. PRE.2017.058).

### 4.3 Gustometer system

This study was conducted using a custom-built liquid delivery system created for use in MRI (36). Liquid volumes of 0.9 mL were delivered over 3 seconds via programmable syringe pumps. The pumps were controlled by task scripts, which were written in Matlab (2016a; Mathworks, USA). Each pump held one of three 50cc syringes, containing either sugar-sweetened lemonade (SU), non-nutritive sweetened lemonade (SW), or tap water, which was connected to a 10-metre length of PVC plastic tubing (Fischer Scientific; 3mm diameter, 0.75mm wall thickness) that were held together in a sleeve. The tubes were affixed to a pacifier, which held them in the participant’s mouth. The plastic tubing and syringes were all commercially manufactured products designed for clinical use. A new set of syringes and tubing were used for each participant and were discarded after single use for hygiene purposes.

### 4.4 Screening - Discrimination task

During the screening session, participants completed a two forced-choice task, in which they received two liquids from the pump system and rated whether the two flavours were the same or different. On each trial, the word ‘TASTE’ appeared for 3s on the screen to indicate the delivery of either SW or SU. Participants were instructed to hold the liquid in their mouth until the word ‘SWALLOW’ appeared on screen, which was also presented for 3s. Then, they indicated whether they thought the two tastes were different or the same. Finally, participants rated how confident they were in their decision on a VAS scale (*0 = Not at all*, *100 = Extremely*). They had 5s to respond to each question, using a mouse. This task consisted of 40 trials, separated by a 500ms inter-trial interval: 10 for congruous SU-SU, 10 for congruous SW-SW, 10 for incongruous SU-SW, 10 for incongruous SW-SU. After every 10 trials, participants received a 1 mL rinse of water.

### 4.4 MRI Acquisition

MRI data were collected on a 3T Siemens Prisma scanner that was fitted with a 32-channel head coil at the Cognition and Brain Sciences Unit in Cambridge, UK. The following parameters were used for echo-planar imaging: repetition time (TR) = 1503 ms, echo time (TE) = 32.40 ms, 2mm isotropic voxels, slice thickness = 2mm, flip angle 74°, field of view (FOV) = 256mm, matrix = 96 x 96, 60 axial slices). We acquired 1660 multiband echo planar 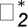-weighted volumes (GE-EPI) with a multi-band acceleration factor of 3 for the probabilistic task. For the deterministic task, 779 volumes were collected using the same multiband parameters. To improve localization of the functional data, a high-resolution T1-weighted anatomical scan was acquired during the same scan session (TR = 2250ms, TE = 3.02ms, 1mm isotropic voxels, slice thickness = 1mm, flip angle = 9°, FOV = 192mm, 192 slices).

### 4.5 Neuroimaging - Probabilistic task

Participants completed an event-related probabilistic conditioning task, consisting of 144 trials (40-minute duration). On each trial, participants were presented with a cue for 1.5s, which included a pie-chart indicating the probability of receiving either SW or SU, and images of beverages on the right and left side of the screen (**Figure 1B**). The probabilities were 50%-50% or 25%-75%, and these were respected on all trials. The cue order and location of SW or SU beverages was pseudorandomized across trials for each participant, resulting in 6 conditions (3 probability levels, plus left and right side placement of the cue). Following the cue, participants were presented with the word ‘TASTE’ and the image of a droplet for 3s, which signalled the delivery of one of the liquids. Participants were instructed to hold the liquid in their mouth until seeing the word ‘SWALLOW’, which was presented for 5s. During this time, participants used a sliding scale, ranging from 1 to 100 with anchors of ‘Sugar’ and ‘Sweetener’, to indicate whether they had received SW or SU (Figure 1). Responses were used to derive both accuracy and confidence measures, where responses >51 were classified as correct and <51 incorrect. Confidence was calculated as the absolute difference of the response from the midpoint, which was divided by the range and multiplied by 100: [|Response - 50|/50]*100. The starting position of the slider was randomised across trials, which ensured that confidence ratings were decoupled from the distance the slider was moved. A fixation cross of variable duration (0.75, 1.12, 1.5, 1.9 or 2.25s, pseudorandomized) was presented between each stimulus. After every 7-11 trials, a rinse was delivered to wash out the lemonade flavour.

In order to refill the pumps, two pauses, lasting approximately 3-5 min, were added to the task. Participants were instructed to look at a fixation cross during the breaks.

### 4.6 Neuroimaging - Deterministic task

The second fMRI task was a deterministic conditioning paradigm that took 25 minutes to complete. During this task, participants were presented cues that indicated which liquid (SU, SW, water) would be delivered (**Figure 1C**). Water delivery was always respected, but on 50% of the SU and SW trials, cues were violated (that is, if the cue indicated SW, SU was delivered, and vice versa). After each cue, the word ‘TASTE’ was shown for 3s, indicating the liquid delivery from the pump. Participants held the liquid in their mouth until the word ‘SWALLOW’ appeared in addition to the question ‘How pleasant was the liquid?’ (5s duration). VAS pleasantness ratings were made using the button box. As in the probabilistic task, each event was followed by a fixation cross of variable duration (0.75, 1.12, 1.5, 1.9 or 2.25s, pseudorandomized).

Participants completed 75 trials, comprising five different conditions of 15 trials each: water delivery, sugar-sweetened lemonade cue and delivery (SU expected), non-nutritive sweetened lemonade cue and delivery (SW expected), sugar-sweetened lemonade cue and non-nutritive sweetened lemonade delivery (SU unexpected, and non-nutritive sweetened lemonade cue and sugar-sweetened lemonade delivery (SW unexpected).

### 4.7 Data analysis – Behaviour

Behavioural data were analysed using linear mixed-effects modelling (LMMs) in R (v3.6.3). For the probabilistic task, we examined the effect of flavour delivered (SU, SW), probability level (unexpected [25%], chance [50%], expected [75%]), and their interaction on participant accuracy and confidence. Specifically, the probability level of a given trial was defined relative to the flavour delivered. That is, if a cue indicated 25% probability of SU and 75% probability of SW, and sugar was delivered, this would be considered unexpected. If the same cue was followed by delivery of the non-nutritive sweetener, this would be considered expected. Probability levels were compared using non-orthogonal contrast coding, which compared both unexpected versus chance and expected versus chance trials. This model included random intercepts for flavour and probability level, which were nested within the participant’s random effect. Results were corrected for multiple comparisons using a Bonferroni correction across the two models (p= 0.05/2 = 0.025).

For the deterministic task, we determined the effect of flavour delivered (SU, SW), violation conditions (expected, unexpected), and their interaction on subjective pleasantness ratings. Random intercepts for flavour type and violation were nested within the subject’s random effect. Because expectancies relating to water trials were not manipulated in either task, they were excluded from behavioural analyses. All data met the assumptions of the aforementioned statistical tests.

### 4.8 Data analysis – Neuroimaging

Functional MRI data were preprocessed and analysed using SPM12 software (Wellcome Department of Cognitive Neurology, London, UK) in MATLAB (MathWorks, Natick, MA). Preprocessing included slice timing correction, within-subject image realignment to the mean volume, voxelwise weighted echo combination (summation based on local 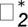 measurements (37)), coregistration of functional images with the □_1_-weighted anatomical scan, spatial normalisation to the Montreal Neurological Institute (MNI) template and spatial smoothing using a 6mm full-width-at-half -maximum Gaussian kernel. The resulting time-series for each functional run were high-pass filtered (1/128Hz), and serial autocorrelations were estimated using a FAST model. Following exclusions for quality control (e.g., excessive head motion, task non-responsiveness), pre-processed data were available from n = 21 and n = 23 participants for the probabilistic and deterministic tasks, respectively.

For both tasks, images were analysed in an event-related manner using a general linear model (GLM), and 6 nuisance regressors were included in the GLM to account for motion-related artefacts (rotation and translation in X, Y, Z planes). The probabilistic task included nine explanatory variables: the cue type (SU 25% & 75% SW, 50% SU & 50% SW, 75% SU & 25 % SW) and the flavour delivered, depending on the probability of receiving it (unexpected [25%], uncertain [50%] and expected [75%] SU delivery; unexpected [25%], uncertain [50%] and expected [75%] SW delivery). Regressors of no interest were included for swallow events, water delivery and refilling of the syringes. Contrasts were generated to assess the effects of cue and flavour type on the BOLD response across differing probability levels. Specifically, we estimated contrasts for expected versus unexpected (e.g., 75% SU versus 25% SU), expected versus uncertain (e.g., 75% SU vs 50% SU), and unexpected versus uncertain (e.g., 25% SU vs 50% SU) SU and SW cues, as well as their delivery. To examine potential differences in brain responses to each flavour, we computed three additional contrasts for: (1) SW versus SU delivery on uncertain trials (50%-50% cues), (2) SU delivery relative to the implicit baseline and (3) SW delivery relative to the implicit baseline. Finally, we examined the parametric effect of confidence on BOLD responses to (1) SU delivery, (2) SW delivery, (3) SU vs SW delivery.

Nine explanatory variables were modelled for the deterministic task: 1) the cue (SW, SU, water); 2) the flavour received across violation conditions (expected SU, unexpected SU, expected SW, unexpected SW, water); and 3) swallow events. Swallow events were considered a nuisance regressor. We included pleasantness ratings as a parametric modulator for SU and SW delivery. To determine the effect of expectation on brain responses to each flavour, we generated four contrasts: expected versus unexpected SU, expected versus unexpected SW, expected SU versus expected SW and unexpected SU versus expected SW. To evaluate brain responses to the delivery of SU and SW, regardless of expectation, we created two additional contrasts to compare flavour delivery to the implicit baseline.

Contrast estimates from the first-level contrasts were entered into a second-level effects analysis, using a one-sample t-test (one-tailed) to compare activation levels in the condition of interest. Second-level analyses were initially conducted within *a priori* ROIs before conducting an exploratory whole-brain analysis. ROIs were either generated or selected using the Wake Forest University Pick Atlas (38) and merged across two masks, one including regions implicated in expectation and the other taste processing (**Figure 2**). The expectation mask was composed of bilateral basal ganglia and ventral frontal regions: orbitofrontal cortex, ventromedial prefrontal cortex, midbrain, putamen, caudate and nucleus accumbens. The taste mask included bilateral regions that have been functionally implicated in flavour perception: insula, anterior cingulate cortex, thalamus, and frontal operculum. The nucleus accumbens ROI was inflated by 2; all other regions were generated without inflation. The midbrain and operculum regions were functionally defined based on prior literature, and all other regions were anatomically defined. For the midbrain, a 15mm spherical region was generated from findings of Murray et al. (39), with central coordinates of MNI_X,Y,Z_ = 0, -15, - 9). The bilateral frontal operculum was estimated from the peak locations reported by Small et al. (40), where an 11mm sphere was centred on the following coordinates for the right and left hemisphere, respectively: MNI_X,Y,Z_ = 54, 15, 12 and MNI_X,Y,Z_ = -54, 15, 1.

A small volume correction was applied to our ROI analyses, and statistical maps from the whole-brain analyses were adjusted for multiple tests using cluster-based correction (cluster-defining threshold of p < .001, FWE-corrected cluster p < .05, one-tailed).

## Author Contributions

HZ, KMJ, PCF and EM designed the experiment. EM, HZ, and KMJ recruited participants and collected the data. EM and KMJ analysed the neuroimaging data, with input from HZ, PCF and MLW. MLW and EM analysed the behavioural data. MLW prepared figures, tables and the initial draft of the manuscript. All authors provided critical feedback on the manuscript.

## Acknowledgements

This work was supported by funding from the Bernard Wolfe Health Neuroscience Fund to P.C.F. and H.Z. and a Wellcome Trust Investigator Award to P.C.F. (Reference No. 206368/Z/17/Z). MLW is supported by a Wellcome Trust Sir Henry Wellcome Postdoctoral Fellowship (224107/Z/21/Z).

