## Supplementary Material for "Expectation modulates the hedonic experiences of and midbrain responses to sweet flavour"

1. **Supplementary Results**

*1.1 Liking and internal state ratings*

Liking and internal state ratings of hunger and thirst were recorded at three time points during the MRI experiment, and linear-mixed effects models were estimated to determine the effects of time point and, when appropriate, flavour type on these ratings. Reverse Helmert contrast coding was used to examine differences between time points, such that the mean of each level (i.e., time point) was compared to the mean of the previous levels.

The effects of flavour type and time on liking were not significant (all p’s > .41). The addition of a flavour-by-time interaction effect did not improve model fit indices (χ^2^_(10)_ = 2.74, p = .25), suggesting that liking ratings for sugar and non-nutritive sweetener remained similar for the duration of the experiment.

Models for internal state ratings also identified a main effect of time point on thirst ratings, which, as expected, generally declined throughout the MRI scan [time point 2 vs. 1: B(SE) = -32.65(7.15), p < .0001); time point 3 vs time point 1 & 2: B(SE) = -18.49(6.20), p = .0047)]. Hunger ratings did not differ significantly across the MRI session (all p’s > .45).

**Table S1.** Exploratory whole-brain responses during the probabilistic conditioning task

| **Contrast** | ***kE*** | **Side** | **Region** | **Peak MNI Coordinates** | | | **Size (voxels)** | **Z-**  **statistic** | **Cluster**  **pFWE** |
| --- | --- | --- | --- | --- | --- | --- | --- | --- | --- |
|  |  |  |  | **X** | **Y** | **Z** |  |  |  |
| 50% > 75% & 25% cue | 129 | L | Superior occipital gyrus | -12 | -90 | 22 | 129 | 4.46 | .044 |
|  |  | L | Lingual gyrus, cerebellum | -18 | -66 | -14 | 199 | 4.35 | .006 |
| SW 50% > SU 50% taste | 279 | L | Superior parietal | -8 | -78 | 52 | 607 | 4.14 | < .001 |
| SW taste > baseline | 214 | R | Cerebellum, brainstem, postcentral gyrus (left), precentral gyrus (left), insula (left) | 12 | -38 | -34 | 7554 | 5.25 | .001 |
|  |  | R | Postcentral gyrus, precentral gyrus, anterior insula, putamen | 60 | -18 | 24 | 5154 | 5.08 | .003 |
|  |  | R | Supplementary motor cortex | 6 | 0 | 62 | 490 | 4.67 | .008 |
|  |  | R | Cerebellum | 16 | -62 | -22 | 214 | 4.55 | .006 |
| Average confidence PM2 | 344 | R | Hippocampus/ parahippocampal gyrus (bilateral), brainstem | 20 | -24 | -12 | 5521 | 5.75 | < .001 |
|  |  | L | Postcentral gyrus | -60 | -18 | 42 | 416 | 4.77 | .013 |
|  |  | R | Postcentral gyrus, precentral gyrus, central operculum | 54 | -6 | 26 | 1352 | 4.49 | <.001 |
| SU confidence PM2 | 398 | L | Precentral gyrus, postcentral gyrus, central operculum, insula (bilateral) | -44 | -14 | 34 | 30,063 | 5.78 | < .001 |

|  |  | R | Supplementary motor cortex (bilateral) | 2 | -2 | 64 | 1002 | 4.53 | < .001 |
| --- | --- | --- | --- | --- | --- | --- | --- | --- | --- |
|  |  | L | Postcentral gyrus | -20 | -30 | 58 | 398 | 4.52 | .014 |
|  |  | R | Postcentral gyrus | 22 | -30 | 58 | 430 | 4.12 | .010 |
| SW confidence PM2 | 1067 |  | Brainstem, parahippocampal gyrus (bilateral) | 4 | -38 | -14 | 1067 | 5.01 | < .001 |
| SU > SW confidence | 15,929 | L | Precuneus, superior parietal lobule, angular gyrus, cuneus, thalamus | -10 | -54 | 16 | 15,929 | 5.18 | .003 |

**Note:** For whole-brain analyses, significant clusters were identified at a cluster-defining threshold of p < .001 and FWE-corrected at p < .05, one-tailed. MNI coordinates represent the peak voxel within each cluster. Z-statistic corresponds to peak value from each cluster. SU = sugar-sweetened beverage, SW = non-nutritive sweetened beverage.

**Table S2.** Exploratory whole**-**brain responses during the deterministic conditioning task

| **Contrast** | ***kE*** | **Side** | **Region(s)** | **Peak MNI Coordinates** | | | **Size (voxels)** | **Z-**  **statistic** | **Cluster**  **pFWE** |
| --- | --- | --- | --- | --- | --- | --- | --- | --- | --- |
|  |  |  |  | **X** | **Y** | **Z** |  |  |  |
| Unexpected SW > Expected SW | 241 | R | Brain stem, cerebellum | 8 | -38 | -36 | 241 | 4.82 | .004 |
| SU taste > baseline | 322 | L | Postcentral gyrus, central operculum, posterior insula | -60 | -6 | 26 | 3,256 | 5.24 | < .001 |
|  |  | R | Supplementary motor area | 2 | 0 | 62 | 322 | 3.95 | < .001 |
|  |  | R | Cerebellum, calcarine cortex, cuneus | 20 | -64 | -20 | 18,599 | 5.15 | .036 |
| SW taste > baseline | 308 | R | Calcarine cortex, cuneus | 10 | -72 | 12 | 13,378 | 5.39 | < .001 |
|  |  | L | Central operculum, precentral gyrus, posterior insula | -60 | -6 | 26 | 6,000 | 5.35 | < .001 |
|  |  | R | Brainstem | 8 | -38 | -42 | 313 | 4.17 | .040 |
|  |  | L | Putamen | -24 | -16 | -4 | 308 | 4.05 | .042 |
| Pleasantness SU | 228 | L | Postcentral gyrus, precentral gyrus, central operculum | -44 | -18 | 36 | 3,501 | 4.95 | < .001 |
|  |  | L | Cerebellum, calcarine cortex | -18 | -60 | -18 | 9,368 | 4.94 | < .001 |
|  |  | R | Postcentral gyrus, precentral gyrus, central operculum | 62 | -14 | 24 | 2,240 | 4.90 | < .001 |
|  |  | L | Transverse temporal gyrus, parietal operculum | -30 | -30 | 18 | 253 | 4.60 | .032 |
|  |  | R | Brainstem | 8 | -38 | -42 | 253 | 4.15 | .041 |
|  |  | R | Supplementary motor cortex | 4 | -2 | 64 | 228 | 3.75 | .045 |
| Pleasantness SU > SW | 327 | R | Inferior occipital gyrus | 46 | -70 | 8 | 327 | 4.19 | .046 |

**Note:** For whole-brain analyses, significant clusters were identified at a cluster-defining threshold of p < .001 and FWE-corrected at p < .05, one-tailed. MNI coordinates represent the peak voxel within each cluster. Z-statistic corresponds to peak value from each cluster. SU = sugar-sweetened beverage, SW = non-nutritive sweetened beverage.

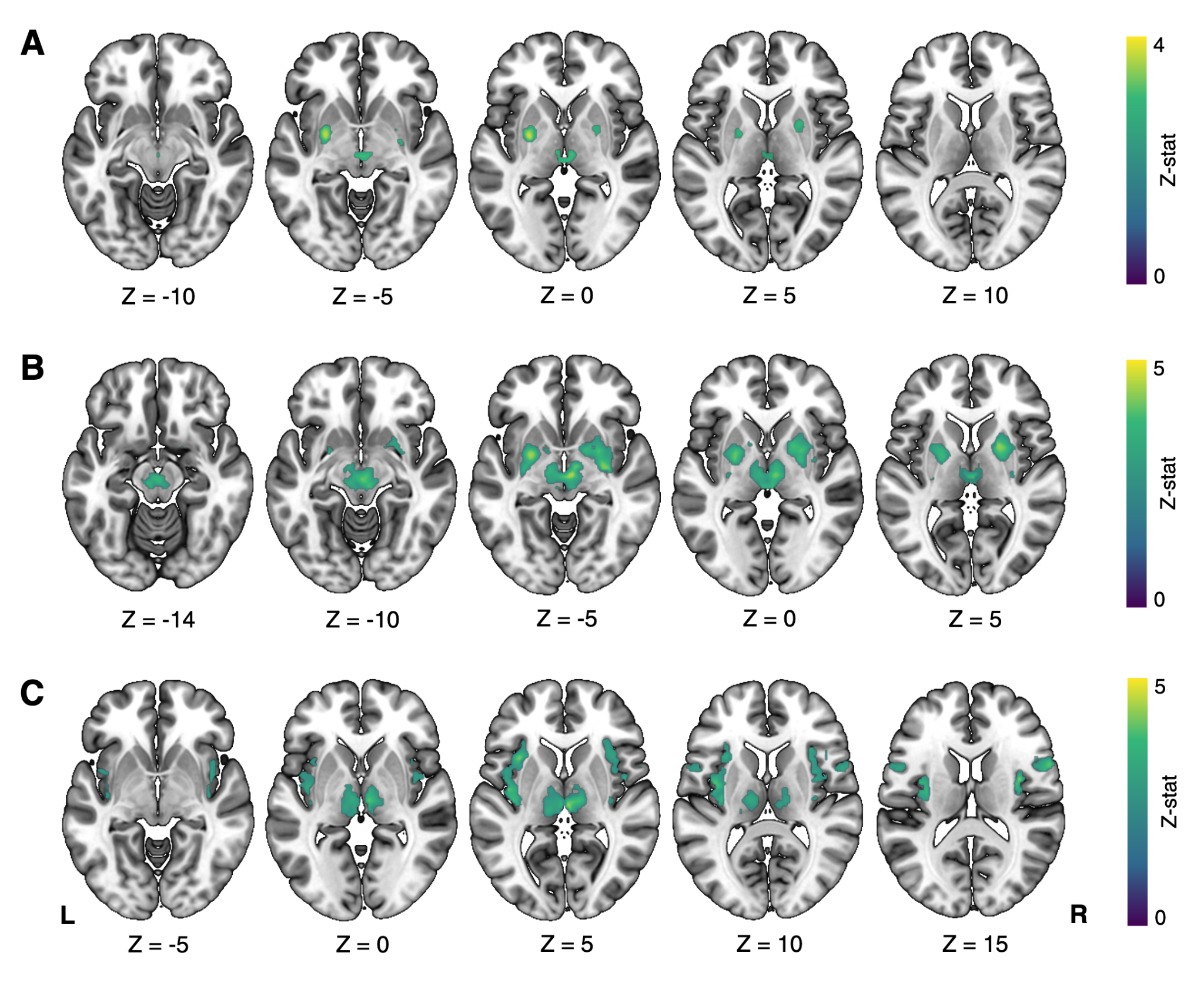

**Figure S1.** *ROI brain responses to sugar and non-nutritive sweetener delivery - Probabilistic task.* **A)** Brain responses to sugar delivery across the expectation mask. **B)** Non-nutritive sweetener delivery increased brain activation in reward and gustatory regions within both expectation and **(C)** taste masks. Clusters within these ROIs were identified using a small-volume correction (voxel-wise p < .001); peak p-values < 0.05 were considered statistically significant. Details on cluster size, coordinates, and associated test statistics, are presented in Table 2.

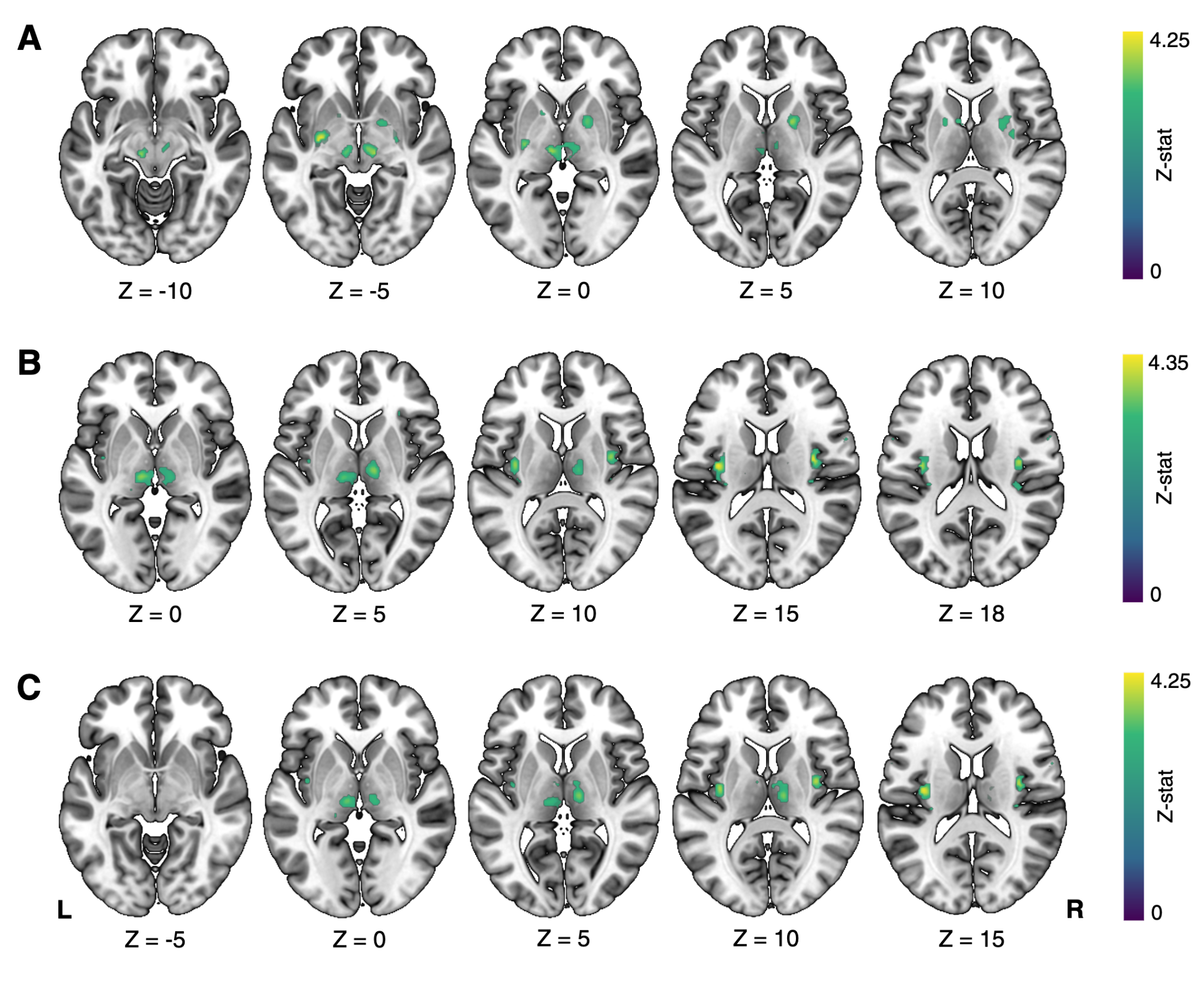

**Figure S2.** *ROI brain responses to sugar and non-nutritive sweetener delivery - Deterministic task.* Brain responses to sugar delivery across the expectation **(A)** and taste **(B)** masks. **(C)** Non-nutritive sweetener delivery elicited similar insular and thalamic activation across the taste mask. Clusters within these ROIs were identified using a small-volume correction (voxel-wise p < .001); peak p-values < 0.05 were considered statistically significant. Details on cluster size, coordinates, and associated test statistics, are presented in Table 4.

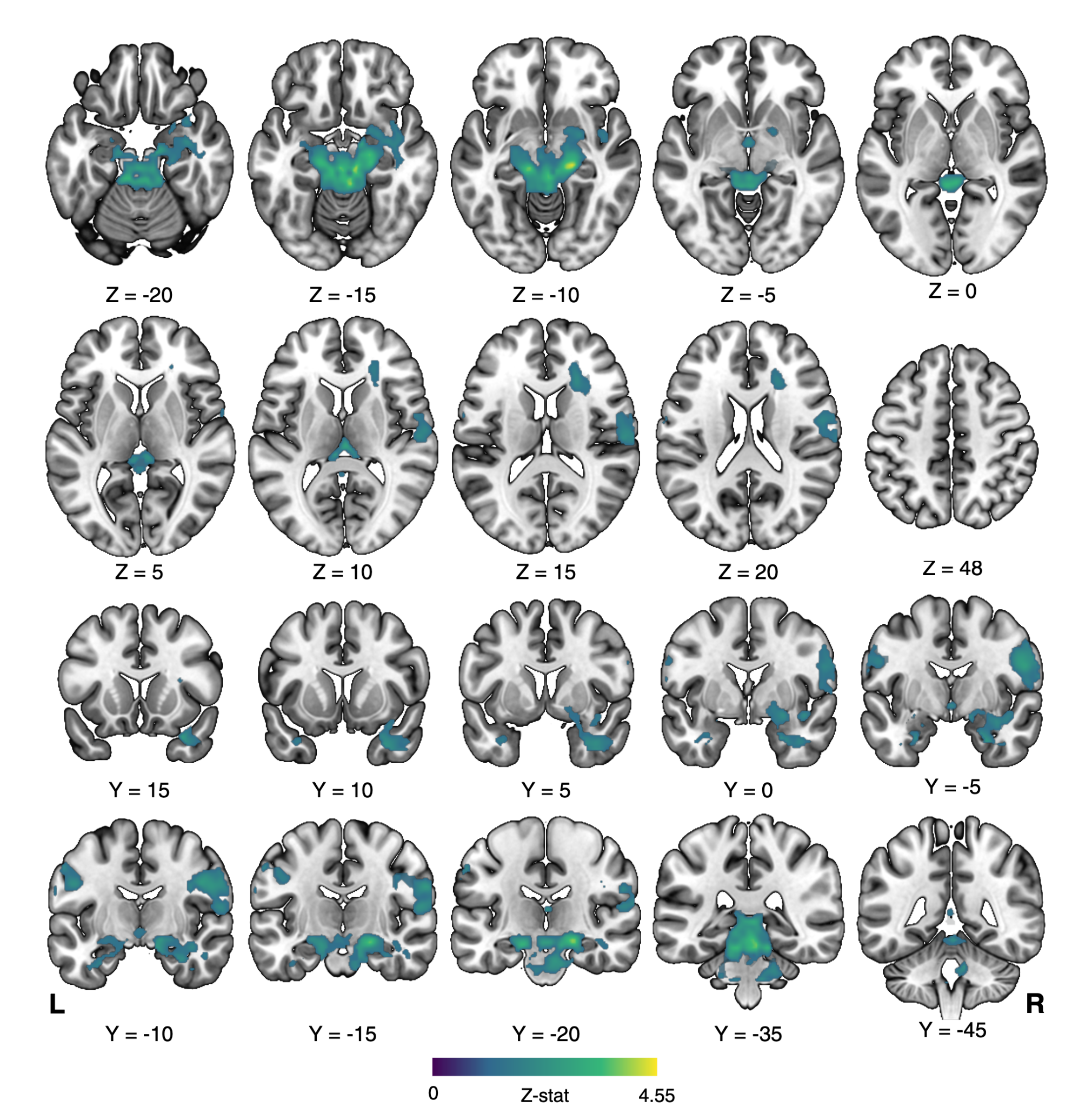

**Figure S3.** *Parametric effect of confidence on brain responses to sweet flavour - whole brain responses.* One-tailed t-test of the parametric effect of confidence on brain responses to both sugar and non-nutritive sweetener delivery. Maps represent significant clusters (voxel-wise p < .001, FWE cluster probability p < .05). For details on cluster size, coordinates, and associated test statistics, see Table 3.

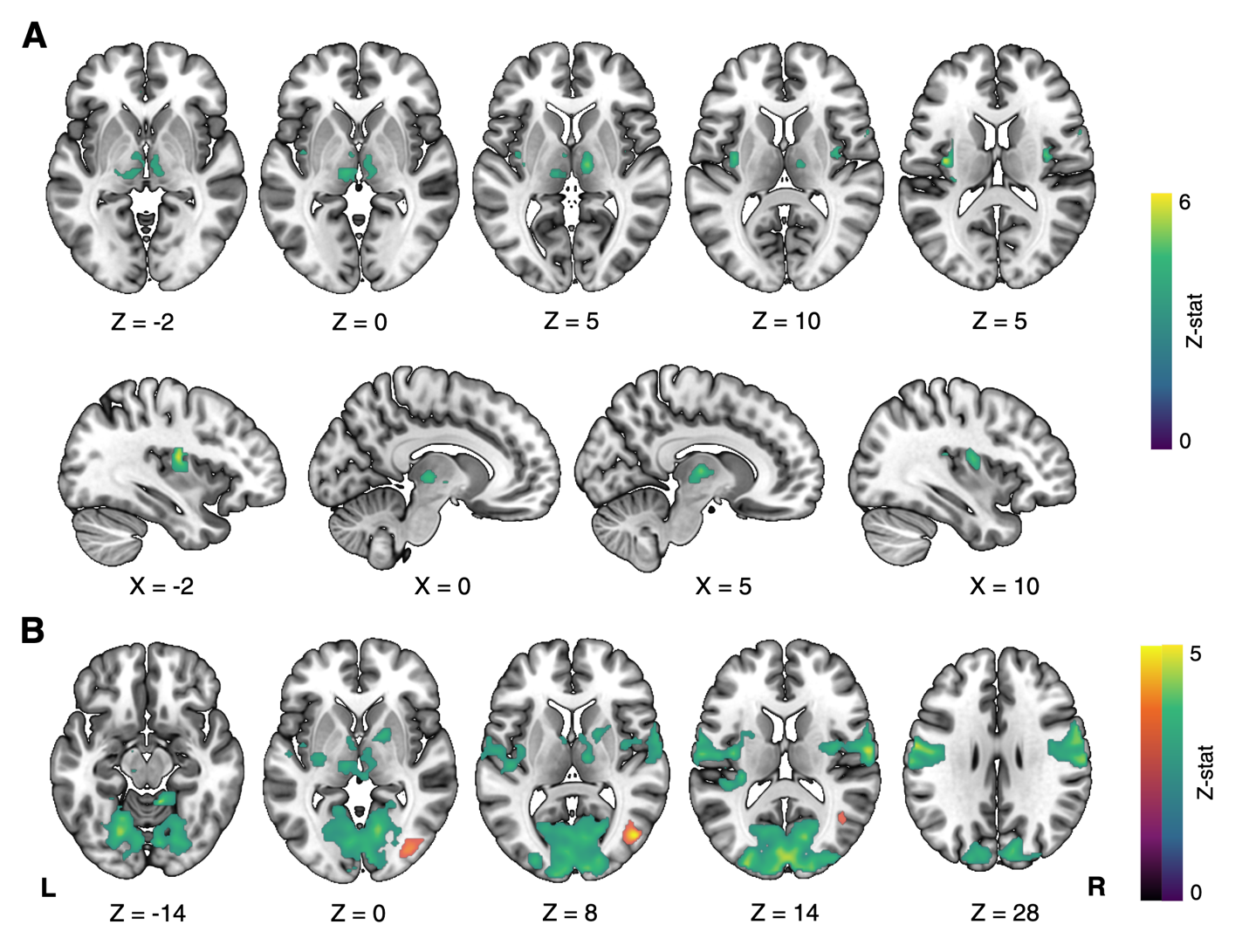

**Figure S4.** *Parametric effect of pleasantness on brain responses to sweet flavour - whole-brain responses.* **(A)** The parametric effect of pleasantness augmented brain responses to sugar-sweetened beverage delivery in across the taste mask during the deterministic task. ROI clusters were identified using a small-volume correction (voxel-wise p < .001); peak p-values < 0.05 were considered statistically significant. **(B)** One-tailed t-test of the parametric effect of pleasantness on brain responses to sugar (blue/green) and sugar versus non-nutritive sweetener delivery (orange). Maps represent significant clusters (voxel-wise p < .001, FWE cluster probability p < .05). For details on cluster size, coordinates, and associated test statistics, see Table S2.
